# Coordination between growth and stress responses by DELLA in the liverwort *Marchantia polymorpha*

**DOI:** 10.1101/2021.02.11.430820

**Authors:** Jorge Hernández-García, Rui Sun, Antonio Serrano-Mislata, Keisuke Inoue, Carlos Vargas-Chávez, David Esteve-Bruna, Vicent Arbona, Shohei Yamaoka, Ryuichi Nishihama, Takayuki Kohchi, Miguel A. Blázquez

**Author notes:** Co-first authors (Jorge Hernández-García, Rui Sun).

## Abstract

Plant survival depends on the optimal use of resources under variable environmental conditions. Among the mechanisms that mediate the balance between growth, differentiation and stress responses, the regulation of transcriptional activity by DELLA proteins stands out. In angiosperms, DELLA accumulation promotes defense against biotic and abiotic stress and represses cell division and expansion, while loss of DELLA function is associated with increased plant size and sensitivity towards stress^1^. Given that DELLA protein stability is dependent on gibberellin (GA) levels^2^, and GA metabolism is influenced by the environment^3^, this pathway is proposed to relay environmental information to the transcriptional programs that regulate growth and stress responses in angiosperms^4,5^. However, *DELLA* genes are also found in bryophytes, whereas canonical GA receptors appeared only in vascular plants^6–10^. Thus, it is not clear whether these regulatory functions of DELLA predated or emerged with typical GA signaling. Here we show that, as in vascular plants, the only DELLA in the liverwort *Marchantia polymorpha* also participates in the regulation of growth and key developmental processes, and promotes the tolerance towards oxidative stress. Moreover, part of these effects is likely caused by the conserved physical interaction with the MpPIF transcription factor. Therefore, we suggest that the role in the coordination of growth and stress responses was already encoded in DELLA from the common ancestor of land plants, and the importance of this function is justified by its conservation over the past 450 M years.

## Results and Discussion

### MpDELLA accumulation affects cell division

The genome of *M. polymorpha* encodes a single Mp*DELLA* gene (Mp5g20660)^7,11^. To investigate its biological function, we generated transgenic plants overexpressing Mp*DELLA* either under the control of the CaMV *35S* promoter with a carboxy-terminal Citrine fusion (*_pro_35S:*Mp*DELLA-Cit*), or the *M. polymorpha ELONGATION FACTOR1α* (Mp*EF1α*) promoter with the 3xFLAG epitope (referred to as _*pro*_Mp*EF:*Mp*DELLA*). In all cases, Mp*DELLA* constitutive overexpressors displayed smaller thallus sizes than the wild type, which was already evident in two-week-old plants (Figures 1A, 1C, and S1A). Introducing additional copies, native-promoter driven translational fusion with the β-glucuronidase (GUS) reporter (*g*Mp*DELLA-GUS*) had a moderate, but similar effect (Figures 1A, 1C, and S1A). As members of the GRAS family transcriptional regulators, DELLA proteins have been shown to function in the nuclei of angiosperms^12,13^. Nuclear localization of MpDELLA was also observed in *g*Mp*DELLA-Cit* plants (Figure S1J). This allowed the construction of Mp*DELLA-GR* lines, which constitutively expressed Mp*DELLA* fused with the rat glucocorticoid receptor. Dexamethasone (DEX)-induced growth impairment was observed (Figure S1C and S1E). These results support that DELLA accumulation inhibits vegetative growth in *M. polymorpha*, similarly to alterations in the sizes of several flowering plant species^14–17^.

**Figure 1.**
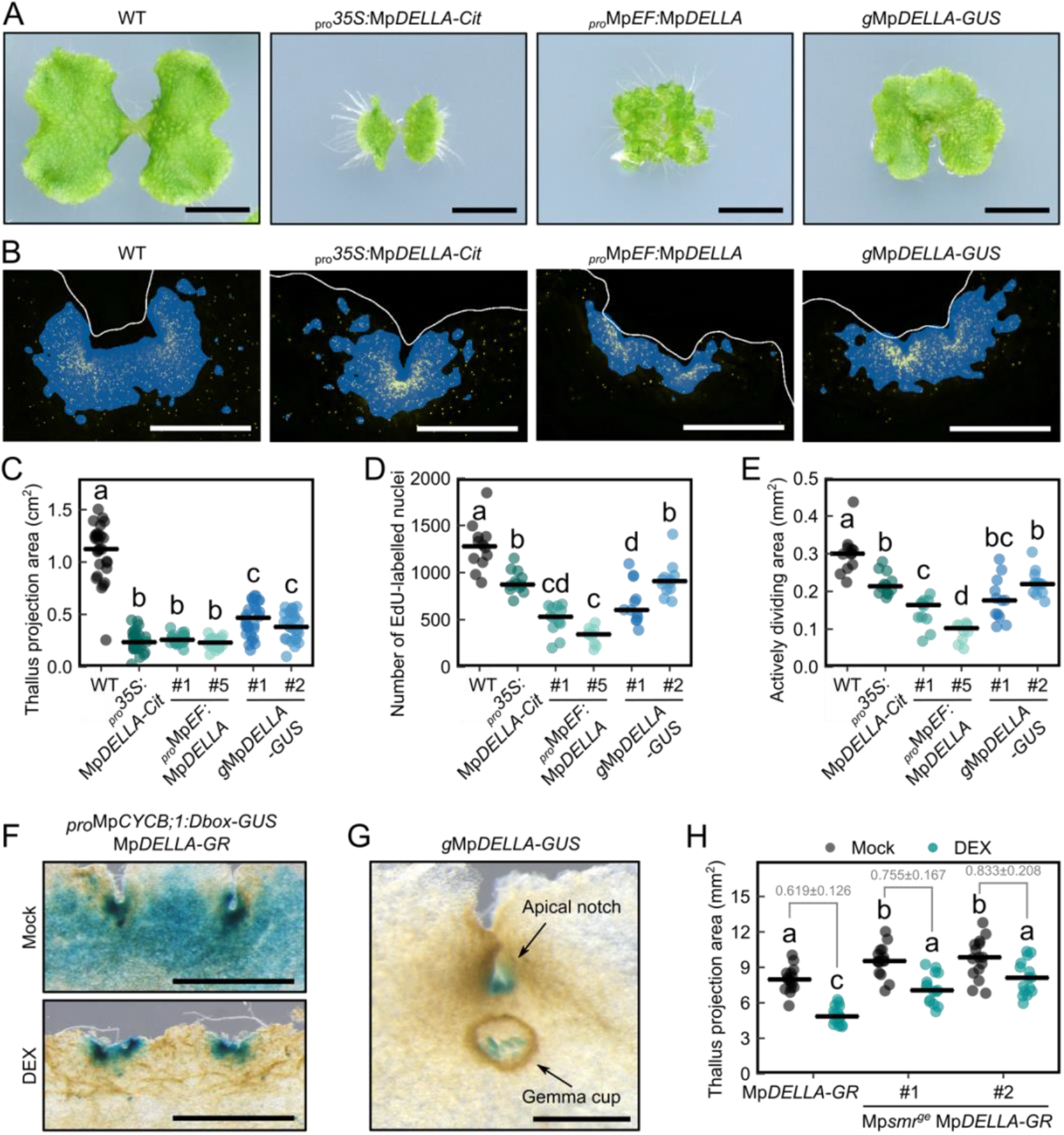
Mp*DELLA* overexpression inhibits plant growth via cell division. See also Figure S1. (A) Morphology of 14-day-old gemmalings in Mp*DELLA* overexpression lines. Scale bar, 5 mm. (B) Apical notches of 7-day-old gemmalings labelled with EdU (yellow signals). Plant boundaries are marked with white lines, and blue color indicates the area occupied by dividing cells (See STAR methods for definition). Scale bar, 500 μm. (C) Measurement of plant sizes in 14-day-old gemmalings. n=26 for *_pro_35S*:Mp*DELLA-Cit*; n=27 for others. (D) Number of EdU-labelled nuclei in the apical notches of 7-day-old gemmalings. n=10 for *_pro_*Mp*EF:*Mp*DELLA* #5; n=12 for others. (E) Quantification of actively dividing areas in the apical notches of 7-day-old gemmalings. n=10 for *_pro_*Mp*EF:*Mp*DELLA* #5; n=12 for others. (F) Images of 9-day-old Mp*DELLA-GR* Mp*CYCB;1-GUS* plants stained for GUS activity after mock or 1 μM DEX treatment for 3 days. Scale bar, 1 mm. (G) A representative image of 21-day-old g*MpDELLA-GUS* plant stained for GUS activity. Scale bar, 500 μm. (H) Measurement of plant sizes in 7-day-old Mp*smr^ge^* Mp*DELLA-GR* gemmalings after mock or 1 μM DEX treatment for 5 days. Ratio of plant sizes (±propagated SE) for each pair is shown in grey. n=15. All plants were grown under continuous white light except long-day conditions in H. In C, D, E, H, dots represent individual plants, and the horizontal lines represent mean values. Statistical groups are determined by Tukey’s Post-Hoc test (p<0.05) following one-way ANOVA.

In Arabidopsis, one of the mechanisms proposed for controlling plant size is the DELLA-dependent reduction of cell proliferation rate^18–20^. To investigate if cell division is affected in Mp*DELLA* overexpressors, we labelled S-phase cells with 5-ethynyl-2’-deoxyuridine (EdU) and observed the nuclear signals around the apical region. All overexpressor lines showed significant reduction in the total number of EdU-positive nuclei, which were distributed in a smaller area compared with wild-type plants (Figures 1B, 1D-E, and S1D, S1F-G). For further confirmation of the cell-cycle regulation by Mp*DELLA*, we introduced a G2-M phase reporter (*_pro_*Mp*CYCB;1:Dbox-GUS*) into the Mp*DELLA-GR* background. After two-day’s treatment with 1 μM DEX, the spatial range of GUS signals was largely restricted compared to the mock group (Figure 1F), suggesting reduction in active cell divisions.

Histochemical analysis of *_pro_MpDELLA:GUS* and *g*Mp*DELLA-GUS* plants showed that MpDELLA is broadly expressed in the thallus, but natively expressed MpDELLA protein preferentially accumulated in the apical notch region, where cell division actively occurs (Figures 1G and S1H). This is comparable with the observations in Arabidopsis that DELLA proteins are expressed in the shoot and root apical meristems^19,21,22^. Similarly, Mp*DELLA* may also restrict growth by inhibiting cell proliferation in the meristematic regions of *M. polymorpha*.

Cyclin-dependent kinase inhibitors (CKIs) have been shown to participate in DELLA-mediated decrease of cell proliferation in Arabidopsis^18,19^. The *M. polymorpha* genome contains two CKI genes, Mp*SMR* (Mp1g14080) and Mp*KRP* (Mp3g00300), which belong to the plant-specific SIAMESE (SIM) protein family and the conserved Kip-related proteins (KRP), respectively^11^. To test genetically if MpDELLA acts through MpSMR to control thallus size, we introduced Mp*SMR* loss-of-function mutations in a Mp*DELLA-GR* line using the CRISPR/Cas9 system^23^ (Figure S1K) and examined growth in the absence and the presence of DEX. Gemmalings carrying the Mp*smr* alleles were moderately larger than the wild type in mock conditions. More importantly, the growth inhibition caused by activation of Mp*DELLA-GR* was attenuated in the Mp*smr^ge^* mutants (Figure 1G), supporting the functional relevance of cell division in Mp*DELLA*-mediated growth restriction. Mp*smr* alleles did not fully abolish the response to DEX induction, indicating that Mp*DELLA* might also suppress cell proliferation through additional pathways.

Taken together, these results suggest that the regulation of plant size through the interference with cell division is a shared DELLA function in land plants.

### MpDELLA regulates development through physical interaction with MpPIF

Distribution of the *g*Mp*DELLA-GUS* signal was also detectable inside gemmae cups, preferentially in developing gemmae (Figures 1H and S1I). Interestingly, both constitutive and induced Mp*DELLA* overexpression exhibited a loss of gemma dormancy, revealed by early gemma germination inside the gemma cups (Figures 2A and 2B). This effect on gemma dormancy resembles the capacity to germinate in the dark of Mp*pif^ko^*, which is a loss-of-function mutant of Mp*PHYTOCHROME INTERACTING FACTOR* (Mp*PIF*, Mp3g17350)^24^. Indeed, we observed a similar loss of dormancy in gemma cups of Mp*pif^ko^* (Figures 2C and S2A). Reciprocally, DEX induction was found to promote gemma germination in darkness for Mp*DELLA-GR* lines (Figure S2B). In addition, Mp*DELLA* overexpressors displayed a significant delay in the induction of sexual reproduction (Figure 2D), which has also been observed in Mp*pif^ko^* ^25^. These similarities indicate a possible functional connection between Mp*DELLA* and Mp*PIF*, which has been previously reported in Arabidopsis for the regulation of apical hook formation and other developmental processes^26–28^.

**Figure 2.**
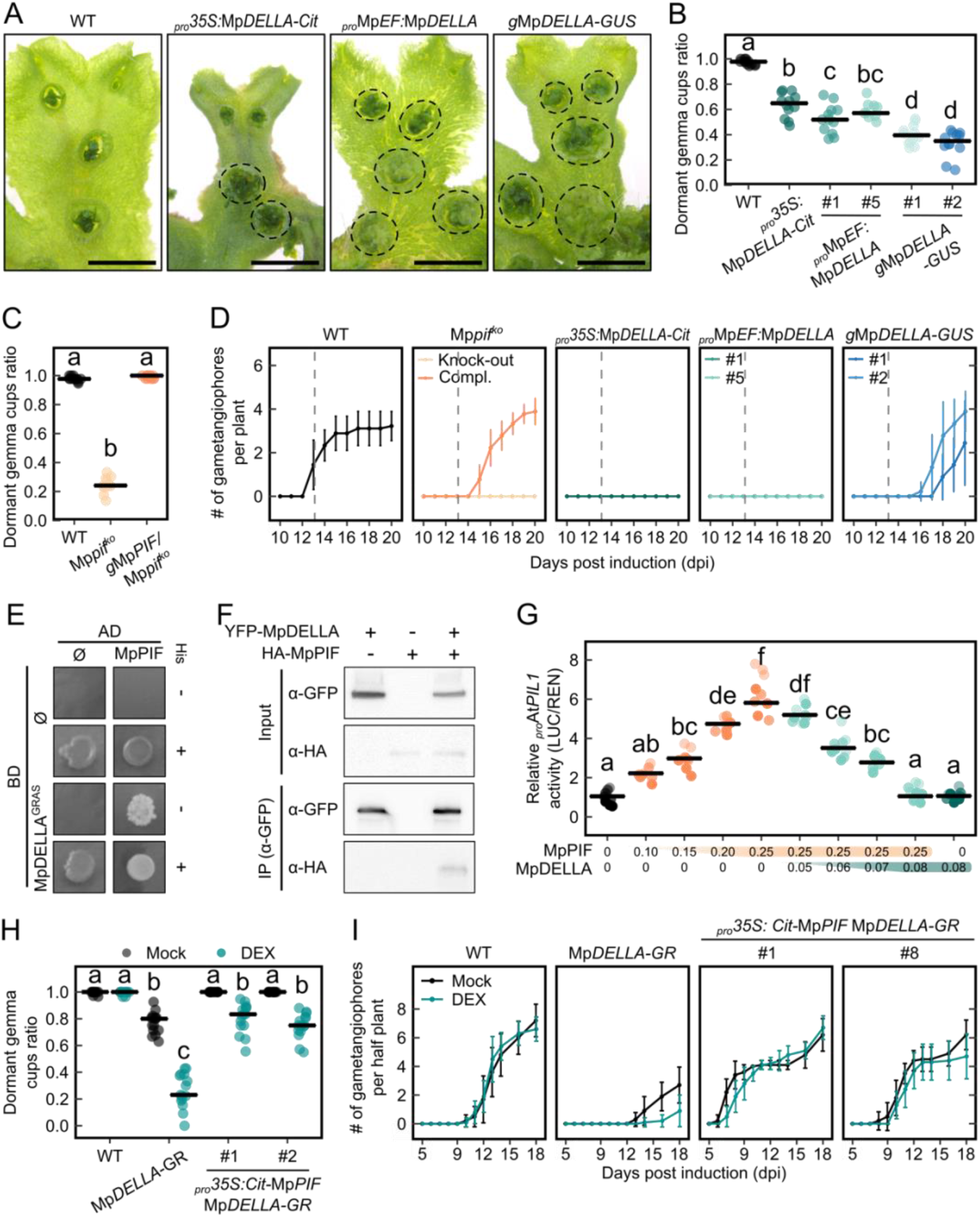
Functional interaction between MpDELLA and MpPIF. See also Figure S2. (A) Images of representative 28-day-old plants showing premature gemma germination inside the cups of Mp*DELLA* overexpression lines. Dashed circles indicate non-dormant gemma cups. Scale bar, 5 mm. (B) Proportion of dormant gemma cups in 28-day-old MpDELLA overexpressors. n=12. (C) Proportion of dormant gemma cups in 28-day-old Mp*pif^ko^* mutants. n=12. (D) Progress of gametangiophore formation in Mp*pif^ko^* mutants and Mp*DELLA* overexpression lines, after induction with far-red light. n=9. (E) Physical interaction between MpDELLA and MpPIF shown by yeast two-hybrid assay. BD and AD denote the fusions to the GAL4 DNA binding domain or the activation domain, respectively (F) Physical interaction between YFP-MpDELLA and HA-MpPIF shown by co-immunoprecipitation after agroinfiltration in *N. benthamiana* leaves. (G) Transient expression assay of the At*PIL1:LUC* reporter in *N. benthamiana* leaves after agroinfiltration with different levels of Mp*PIF* and Mp*DELLA* (shown below the graph as infiltrated OD_600_). n=9 in total. (H) Quantification of gemma cup dormancy in 30-day-old *_pro_*35S:Mp*PIF-Cit* Mp*DELLA-GR* plants, after treatment with mock or 1 μM DEX for 20 days. (I) Progress of gametangiophore formation in *_pro_35S*:Mp*PIF-Cit* Mp*DELLA-GR* plants, after induction with far-red light and treatment with mock or 1 nM DEX. n=10. In A, B and C, plants were grown on ½ Gamborg’s B5 plates with 1% sucrose under cW. In B, C, and H, dots represent individual plants. In G dots represent biological replicates from three independently performed experiments. All horizontal lines represent total mean values. Statistical groups are determined by Tukey’s Post-Hoc test (p<0.05) following ANOVA analysis. In D and I, error bars represent standard deviation.

In Arabidopsis, DELLA proteins interact physically with the PIF transcription factors and prevent their binding to downstream targets^27,28^. It is likely that this mechanism is also conserved in *M. polymorpha*, since we observed that MpDELLA interacts physically with MpPIF in a yeast two-hybrid assay, *in vivo* by co-immunoprecipitation, and also by Bimolecular Fluorescence Complementation assays (Figures 2E, 2F and S2C). Further MpPIF deletion analyses suggested that the GRAS domain of the MpDELLA protein specifically interacts with the bHLH domain of MpPIF (Figures S2D and S2E), paralleling results seen in Arabidopsis^27^. Inhibitory effect of the interaction was verified by dual-luciferase transactivation assays in tobacco. In a dose-dependent manner, MpDELLA inhibited the MpPIF-activation of the At*PIL1* promoter (Figure 2G), a known direct target for PIFs in Arabidopsis^29^.

To assess the biological relevance of the interaction between MpDELLA and MpPIF, we tested if an increase in the dosage of MpPIF would suppress the phenotypical defects caused by high MpDELLA levels. Indeed, the reduction of gemma dormancy in gemma cups caused by MpDELLA induction was notably attenuated in *_pro_35S*:*Cit-*Mp*PIF* Mp*DELLA-GR* plants (Figures 2H and S1A-B). Similarly, gemma germination in darkness and the delay in gametangiophore formation of Mp*DELLA-GR* plants were significantly suppressed in the presence of higher Mp*PIF* levels, both in the absence and presence of DEX treatments (Figures 2I and S2F). No rescue of plant growth by elevated Mp*PIF* levels was observed in the double overexpression lines (Figure S2G). Given the normal vegetative growth of Mp*pif^ko^* ^24^, the cell-cycle-repressing function of MpDELLA does not appear to be mediated by MpPIF.

These results suggest that the modulation of developmental programs by DELLA through its interaction with PIF could be a conserved mechanism already present in the common ancestor of land plants. Besides, since Mp*PIF* is a central regulator in red/far-red light signaling, gemma germination and gametangiophore formation are also developmental decisions in response to light conditions for *M. polymorpha*^24,25^. The additional layer of Mp*DELLA* regulation may also indicate an integrated control of growth, development and response to environmental cues, which has been conserved during land plant evolution. However, the actual processes regulated by DELLA-PIF interaction would depend on the genetic context of particular species.

### MpDELLA promotes flavonoid accumulation and tolerance towards oxidative stress

To investigate the downstream targets of the MpDELLA-MpPIF module, we analyzed the transcriptomic changes in *_pro_35S:*Mp*DELLA-Cit* and the Mp*pif^ko^* mutant. As MpPIF proteins are stabilized by far-red light^24^, Mp*pif^ko^* plants were evaluated at 0, 1, or 4 hours after far-red light irradiation (see STAR Methods). Mp*DELLA* overexpression caused the upregulation of 1483 genes and the downregulation of 560 genes (Figure 3A and Data S1). The analysis of differential gene expression in the Mp*pif^ko^* mutant yielded a total of 339 and 333 genes, up- and down-regulated by at least one time point, respectively (Data S1). As expected, the most abundant set of Mp*PIF*-regulated genes was obtained after the 4-hour far-red treatment (Figure S3A), so we used this set for further analyses. More than half of the upregulated genes in Mp*pif^ko^* were also upregulated in *_pro_35S*:Mp*DELLA*, and there was a statistically significant overlap also among genes downregulated in both genotypes (Figure 3A), indicating a strong correlation between gain of Mp*DELLA*, and loss of Mp*PIF* functions. As a confirmation, three differentially expressed genes were tested by qPCR, and they all showed expression changes consistent with the RNA-seq (Figure S3B).

**Figure 3.**
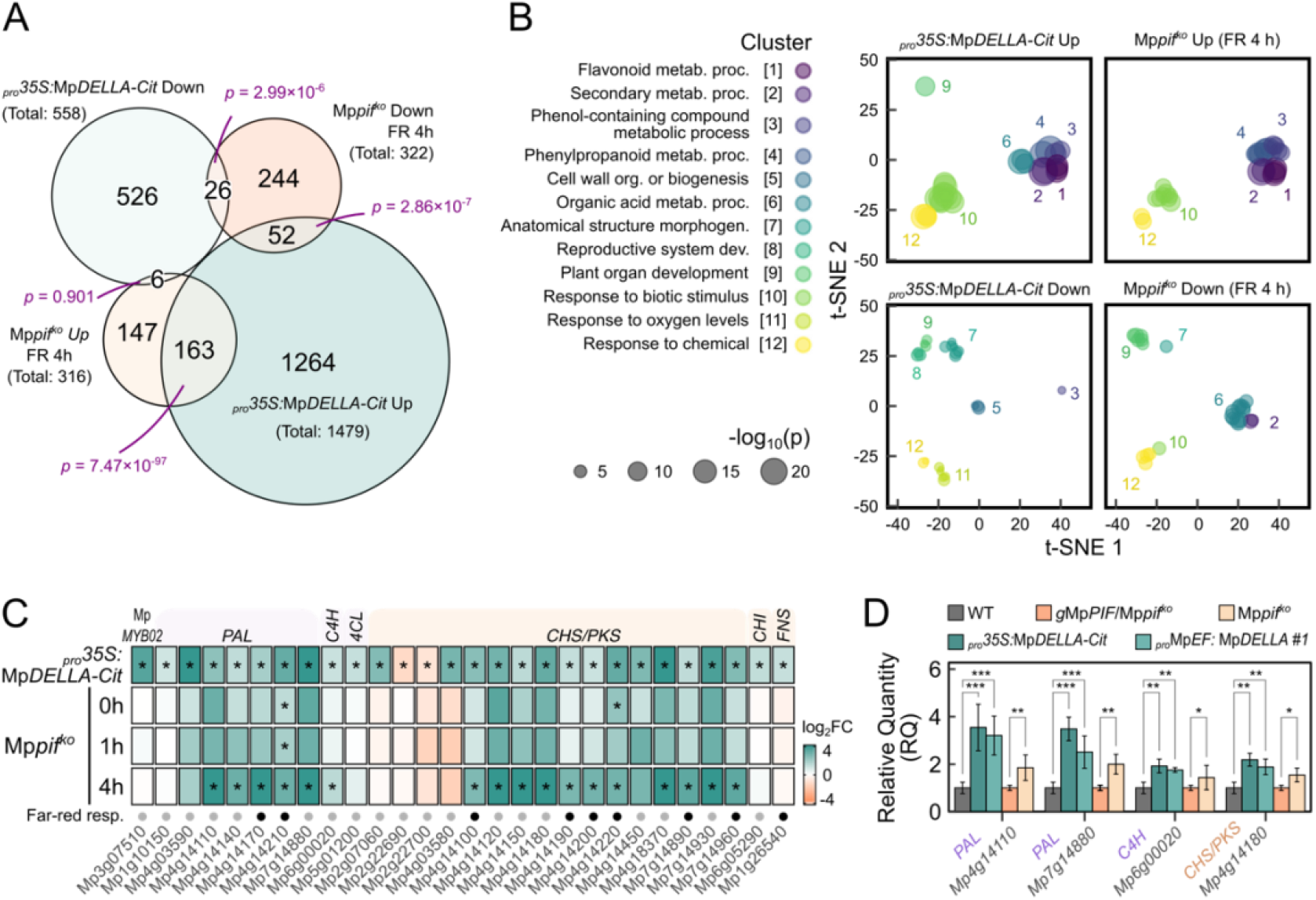
Genome-wide co-regulation of gene expression by MpPIF and MpDELLA. See also Figure S3 and Data S1. (A) Venn diagram showing the overlap between genes differentially expressed in *_pro_35S*:Mp*DELLA-Cit* and between the Mp*pif^ko^* mutant and wild type after 4 hours of far-red light (FR) irradiation. P-values were calculated by Fisher’s exact tests. (B) Two-dimensional t-SNE plot visualizing GO categories over-represented in the sets of genes differentially expressed in *_pro_35S*:Mp*DELLA-Cit* and in Mp*pif^ko^*. (C) Heatmap showing fold changes on log2 scale (log2FC) in the expression of genes encoding enzymes of the flavonoid biosynthesis pathway. Asterisks indicate genes considered as significantly changed (with |log2FC|>1 and adjusted p<0.01 calculated by DESeq2). Black dots in the bottom row indicate genes significantly changed in response to FR irradiation at any time point in either WT or Mp*pif^ko^*. (D) Expression level of selected genes of the flavonoid biosynthesis pathway determined by RT-qPCR. Error bars represent standard deviation of three biological replicates. Asterisks indicate statistically significant differences with respect to the wild type (*, p<0.05; **, p<0.01, after a Student’s t-test).

Gene ontology enrichment analysis highlighted the regulation of stress response and secondary metabolism processes in both *_pro_35S:*Mp*DELLA-Cit* and Mp*pif^ko^* up-regulated datasets, especially with the enrichment of terms involving phenylpropanoid and flavonoid biosynthesis (Figure 3B and Data S1). In particular, many genes encoding PHENYLALANINE AMMONIA LYASE (PAL), CINNAMATE 4-HYDROXYLASE (C4H) or CHALCONE SYNTHASE (CHS) were indeed upregulated, as also confirmed by qPCR (Figures 3C and 3D). In the case of Mp*pif^ko^*, the observed net upregulation of flavonoid biosynthesis genes was mainly due to far-red-induced downregulation in the wild type (Figure S3C), suggesting a suppressive role of MpPIF.

Staining with diphenylboric acid 2-aminoethyl ester confirmed the increased accumulation of flavonoid compounds caused by Mp*DELLA* overexpression or Mp*PIF* loss-of-function (Figures 4A and S4A). Furthermore, Mp*DELLA*-induced increase of flavonoid signals were less evident when Mp*PIF* is also overexpressed in Mp*DELLA-GR* (Figure S4B). Quantitative targeted analysis of flavonoids showed between 1.8 and 7.9 fold induction of luteolin 7-glucuronide, 4,7-dihydroxyflavan-3-ol, and hydroxyl-gouiboutinidol in *_pro_35S:*Mp*DELLA-Cit* (Figure S4C and Table S1). Similar to other plants, increased flavonoid biosynthesis is shown as a protective response against UV-B induced oxidative stress in *M. polymorpha*^30^. The ability to enhance the production of these antioxidant compounds suggests a general function for MpDELLA in stress response, which might be fulfilled in coordination with its inhibition of MpPIF.

**Figure 4.**
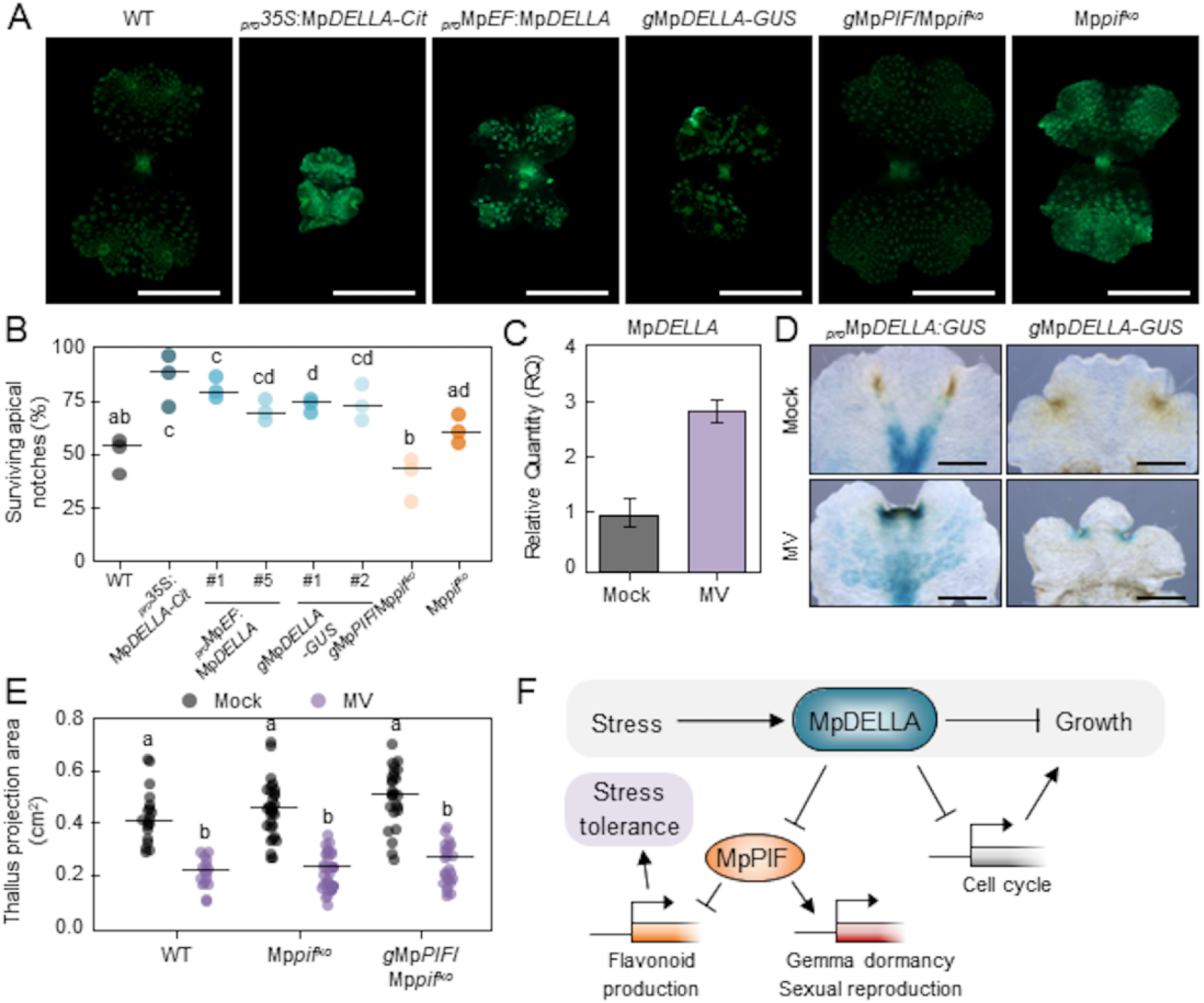
Involvement of Mp*DELLA* in the response to oxidative stress. See also Figure S4 and Table S1. (A) Images of 14-day-old gemmalings, stained with diphenylboric acid 2-aminoethyl ester. More intense fluorescent signals denote higher general flavonoid content. Scale bar, 5 mm. (B) Percentage of surviving apical notches after a 10-day treatment with 100μM MV in different transgenic lines. Data from six independent experiments for Mp*pif^ko^* and gMpPIF/Mp*pif*^ko^, three for others. (C) Relative expression level of Mp*DELLA* determined by RT-qPCR in 14-day-old gemmalings grown in mock or 10 μM MV-supplemented media. Error bars represent standard error from three biological replicates. (D) GUS stained *_pro_*Mp*DELLA*:*GUS* and *g*Mp*DELLA*-*GUS* lines, showing the increased signals in the apical notches of 13-day-old plants after a 6-day treatment with mock or 10 μM MV. (E) Size of 14-day-old gemmalings grown on mock or 0.5 μM MV-supplemented medium. n=19 (WT Mock), 16 (WT MV), 36 (Mp*pif^ko^* Mock), 28 (Mp*pif^ko^* MV), 25 (*g*Mp*PIF*/Mp*pif^ko^* Mock), 24 (*g*Mp*PIF*/Mp*pif^ko^* MV). (F) Model for the regulation of growth, development and stress responses by Mp*DELLA*. Under stress, MpDELLA would accumulate in apical notches protecting them through the MpPIF-dependent production of antioxidant compounds, and suppressing growth by inhibiting cell divisions. The interaction with MpPIF also causes alterations in developmental processes, such as gemma dormancy or gametangiophore formation. All plants were grown under long-day conditions. In B, E, dots represent biological replicates, and the horizontal lines represent mean values. Statistical groups were determined by Tukey’s Post-Hoc test (p<0.05) following ANOVA analysis.

To test if MpDELLA levels influence the response to oxidative stress, we examined the tolerance of plants overexpressing Mp*DELLA* to methyl viologen (MV), an inducer of oxidative stress^31^. Six-day-old gemmalings were transferred to plates containing 100 μM MV for 10 days, after which they were allowed to recover for at least one week. All Mp*DELLA* overexpressing plants, including those with the native Mp*DELLA* promoter, showed a significantly higher survival rate compared to wild-type plants (Figure 4B). These results suggest that the increased production of flavonoids caused by higher Mp*DELLA* levels could be responsible for the protection against oxidative stress. This is further supported by the observation that Mp*pif^ko^* mutants also displayed enhanced resistance to MV (Figure 4B), and that the Mp*DELLA*-dependent tolerance was attenuated by Mp*PIF* overexpression (Figure S4D).

In Arabidopsis, the role of DELLAs in the coordination between growth and stress responses is visualized by GA reduction and accumulation of DELLA in response to certain types of stress, coupled to increased tolerance and a variable degree of growth impairment^32^. Although *M. polymorpha* does not possess a GID1-like GA receptor that might modulate MpDELLA protein stability, we found that exposure of 10-day-old gemmalings to MV provoked an increase in Mp*DELLA* gene expression (Figure 4C and 4D). Such MpDELLA accumulation was concomitant with marked growth arrests and reduction in cell division (Figures 4E and S4E). Interestingly, Mp*pif^ko^* mutants were as large as wild-type plants both in the absence and in the presence of MV (Figure 4E), confirming that the control of *M. polymorpha* thallus size is largely independent of Mp*PIF*.

In summary, our results show that MpDELLA can modulate cell divisions, developmental responses and tolerance to oxidative stress in *M. polymorpha*, through molecular mechanisms that are shared with angiosperms (Figure 4F). This conservation implies that the role in the optimization of growth and the responses to disadvantageous environments would already be encoded in the ancestral land plant DELLA protein, and the canonical GA signaling might have simply hijacked these functions when the pathway emerged in vascular plants.

## Supporting information

DataS1

## Acknowledgements

We thank Eri Nakamura for the pENTR1A-proMpCYCB;1-Dbox plasmid; Shuji Shigenobu and Katsushi Yamaguchi in National Institute for Basic Biology for the help in RNA-seq of Mp*pif^ko^*; Roberto Solano, Maite Sanmartín and all members of the Plant Plasticity Lab for discussions and insightful comments on the manuscript. Work in the authors laboratories has been funded by the Spanish Agencia Estatal de Investigación and FEDER (grants BFU2016-80621-P and PID2019-110717GB to M.A.B) and JSPS/MEXT KAKENHI (JP17H07424 and JP19H05675 to T.K.; 20H04884 to R.N.). J.H.-G. and A.S.-M. were supported by a predoctoral fellowship from the Spanish Spanish Ministry of Education, Culture and Sport (FPU15/01756), and an MSCA Individual Fellowship (H2020-MSCA-IF-2016-746396), respectively. R.S. was supported by the Japanese Government (MEXT) Scholarship Program.

## Author Contributions

M.A.B. and T.K. conceptualized the research. J.H.-G. and R.S. performed the experiments. A.S.-M. performed the genome editing for Mp*SMR*. D.E.-B. did Co-IPs. K.I. did the RNA-seq for Mp*pif^ko^*. R.S. and C.V.-C. did the bioinformatic analysis. V.A. performed flavonoid quantifications. All authors analyzed and/or interpreted the data. M.A.B, J.H.-G. and R.S. wrote the manuscript with the help and corrections from other authors.

## STAR Methods

### RESOURCE AVAILABILITY

#### Lead Contact

Further information and requests for resources and reagents should be directed to and will be fulfilled by the Lead Contact, Miguel A. Blázquez.

#### Materials availability

Plasmids and plant materials generated in this research are all available upon request. Please note that the distribution of transgenic plants will be governed by material transfer agreements (MTAs) and will be dependent on appropriate import permits acquired by the receiver.

#### Data and Code Availability

Raw RNA sequencing datasets generated during this study were deposited to the Short Read Archive at the National Center for Biotechnology Information (NCBI) or the Sequence Read Archive at DNA Data Bank of Japan (DDBJ) under Bioprojects PRJNA695248 and PRJDB11176. The modified ITCN plugin for ImageJ is available at https://github.com/PMB-KU/CountNuclei. R scripts used for processing EdU data, Blast2GO annotation and RNA-seq analysis were deposited to https://github.com/dorrenasun/MpDELLA.

### EXPERIMENTAL MODEL AND SUBJECT DETAILS

#### Plant Materials and Growth Conditions

*Marchantia polymorpha* accession Takaragaike-1 (Tak-1; male)^33^ was used in this study as the wild-type (WT). Female lines Mp*pif^ko^* and *g*Mp*PIF/*Mp*pif^ko^* were previously described as *pif^KO^* #1 and *_pro_PIF:PIF*/*pif^KO^* #1, respectively^24^. *M. polymorpha* plants were cultured on half-strength Gamborg’s B5^34^ medium with 1% agar at 21-22°C. Light conditions are specified in each figure; generally, long day (LD) conditions refer to cycles with 16 hours of light (90-100 μmol m^−2^ s^−1^) and 8 hours of darkness, while continuous white light (cW) was supplemented at the intensity of 50-60 μmol m^−2^ s^−1^.

### METHOD DETAILS

#### Cloning and generation of transgenic *M. polymorpha* plants

Various Gateway-compatible entry vectors related to Mp*DELLA* were generated. The full length CDS, GRAS domain (amino acids 173 to 560), promoter (4.3kb upstream of ATG), and genomic (promoter and CDS) regions were amplified from genomic DNA by PCR using Phusion High-Fidelity Polymerase (Thermo Fisher Scientific) with attB-containing primers, and introduced into pDONR221 (Thermo Fisher Scientific) vector using Gateway BP Clonase II Enzyme mix (Thermo Fisher Scientific) to generate pENTR221-MpDELLA, - MpDELLA^GRAS^, -_pro_MpDELLA and -gMpDELLA, respectively. The CDS region was also amplified with KOD FX Neo DNA polymerase (Toyobo Life Science) and directionally cloned into pENTR/D-TOPO (Thermo Fisher Scientific) to create pENTR-MpDELLA. For pENTR1A-_pro_MpDELLA-short, a slightly shorter promoter region was amplified with PrimeSTAR GXL DNA Polymerase (TaKaRa Bio) and inserted between the *Sal*I and *Not*I sites of pENTR1A (Thermo Fisher Scientific) with T4 DNA ligase (TaKaRa Bio). pENTR1A-gMpDELLA-short was further created by seamless integration of the CDS fragment with the In-Fusion Cloning kit (TaKaRa Bio). Finally both constructs were extended at the 5’ end by In-Fusion insertion to create pENTR1A-_pro_MpDELLA and pENTR1A-gMpDELLA, matching with the lengths of pENTR221 counterparts. To create the vectors for Mp*DELLA* overexpression, pENTR221-MpDELLA and pENTR211-gMpDELLA were recombined with pMpGWB106 and pMpGWB107^35^ using Gateway LR Clonase II Enzyme mix (Thermo Fisher Scientific) to generate pMpGWB106-MpDELLA and pMpGWB107-gMpDELLA, respectively. pENTR-MpDELLA was recombined with pMpGWB310 and pMpGWB313 for the generation of pMpGWB310-MpDELLA and pMpGWB313-MpDELLA, while pENTR1A-_pro_MpDELLA and pENTR1A-gMpDELLA were recombined with pMpGWB304 to generate pMpGWB304-_pro_MpDELLA and pMpGWB304-gMpDELLA. All these binary vectors were introduced into Tak-1 plants.

To monitor the cell division activity, the promoter (3.8 kb upstream of ATG) and coding sequence of the first 116 amino acids (including the destruction box) of Mp*CYCB;1* (Mp5g10030) was amplified with KOD -Plus- Ver.2 (Toyobo Life Science) and ligated into the the *Sal*I and *Eco*RV sites of pENTR1A (Thermo Fisher Scientific) with Ligation high Ver.2 (Toyobo Life Science). The resulting plasmid was recombined with pMpGWB104 and then transformed into the *M. polymorpha* transgenic line Mp*DELLA-GR* #5.

CRISPR/Cas9-based genome editing of Mp*SMR* (*Mp1g14080*) was performed as previously described^36^. The guide RNA was designed upstream of the CDKI domain with Benchling^37^. Double stranded DNA corresponding to the guide RNA protospacers were generated by annealing complementary oligonucleotides and inserted into BsaI-digested pMpGE_En03^36^ by ligation using DNA T4 ligase (Promega), and then transferred to the binary vector pMpGE010 ^36^ using Gateway LR Clonase II Enzyme mix (Thermo Fisher Scientific). *M. polymorpha* transformation was carried out in the transgenic line Mp*DELLA-GR* #5, and targeted loci were examined by sequencing from crude G1 DNA samples and confirmed in G2 plants.

For the construction of Mp*PIF*-Mp*DELLA* double overexpressors, the Mp*PIF* (*Mp3g17350*) coding region containing the stop codon was amplified from cDNA, cloned into pENTR/D-TOPO (Thermo Fisher Scientific), then recombined with pMpGWB105 using Gateway LR Clonase II Enzyme mix (Thermo Fisher Scientific). The resulting construct was transformed into the *M. polymorpha* transgenic line Mp*DELLA-GR* #5.

All the *M. polymorpha* transgenic lines are listed in the Key Resources Table. Transformants were obtained by agrobacterium-mediated transformation from regenerating thalli, using *Agrobacterium tumefaciens* strains GV3101 (pMP90 C58) or GV2260 ^38^.

#### Plant growth and EdU analysis

For the measurement of plant sizes, images of the whole culturing plates were taken vertically above with a digital camera (Canon EOS Kiss X7i). The thallus projection areas were analyzed with ImageJ 1.52a^39^ by thresholding the images with the default algorithm on the blue color channel and batch-measured with the function “Analyze Particles”.

For the detection of S-phase cells, constitutive- and native-promoter MpDELLA overexpressors were grown from gemmae for seven days under cW. Mp*DELLA-GR* lines were grown from gemmae under cW for five days, then transferred onto the plates containing mock solvent or 1 μM dexamethasone (DEX) and cultured for two days. All the plants were labeled with 20 μM 5-ethynyl-2’-deoxyuridine (EdU) in liquid half-strength Gamborg’s B5 medium under cW for 2 h. Then they were fixed with 3.7% formaldehyde for 1 h, washed for 5 min twice with phosphate buffer saline (PBS), and permeabilized in 0.5% Triton X-100 in PBS for 20 min. After two 5-min washes with 3% bovine serum albumin (BSA) in PBS, samples were incubated with the reaction mixture from Click-iT EdU Imaging Kit with Alexa Fluor 488 (Invitrogen) in the dark for 1 h. After staining, samples were protected from light, washed twice with 3% BSA in PBS and soaked in ClearSee solution^40^ for 3-7 days. After that, the samples were mounted to slides with 50% glycerol and observed with Keyence BZ-X700 all-in-one fluorescence microscope.

Z-stacks of fluorescence images were taken in 2-μm steps with the YFP filter (Keyence 49003-UF1-BLA, excitation at 490-510 nm, detection range 520-550 nm) and merged together with the BZ-X Analyzer software (1.3.1.1). EdU-labelled nuclei were marked and counted with a modified version of the ITCN plugin^41^ in ImageJ 1.52a^39^. Spatial coordinates for the nuclei were exported and processed with R scripts^42^ to calculate density maps using the *spatstat* package^43^. Actively dividing area was measured as with nucleus densities higher than 0.001 μm^−2^. See Key Resources Table for the depository of plugins and scripts used.

#### Microscopy & histochemical analysis

For GUS activity assay, plants were vacuum-infiltrated with GUS staining solution (50 mM sodium phosphate buffer pH 7.2, 0.5 mM potassium-ferrocyanide, 0.5 mM potassium-ferricyanide, 10 mM EDTA, 0.01% Triton X-100 and 1 mM 5-bromo-4-chloro-3-indolyl-β-D-glucuronic acid) for 15 min, and then incubated at 37 °C overnight (>16 hours). Samples were de-stained with 70% ethanol and imaged under an Olympus SZX16 stereoscope. To prepare agar sections, stained samples were embedded in 6% agar and sectioned into 100-μm slices with LinearSlicer PRO 7 (DOSAKA EM, Kyoto, Japan), then imaged with Keyence BZ-X700 microscope in the bright-field.

Confocal laser scanning microscopy on *g*Mp*DELLA-Cit* gemma was performed using a Leica TCS SP8 equipped with HyD detectors. A white light laser was used to visualize Citrine (excitation 509 nm). Diphenylborinic acid 2-aminoethyl ester (DPBA) staining was used to visualize flavonoids as previously described^44^. Whole thalli were stained for 15 minutes at 0.25% (w/v) DPBA and 0.1% (v/v) Triton X-100. Epifluorescence microscopy of stained flavonoids in gemmalings was performed on a Leica DMS1000 dissecting microscope using a GFP filter for detection of DPBA fluorescence.

#### Scoring of gemma cup dormancy

To score the dormancy of gemma cups, constitutive- and native-promoter Mp*DELLA* overexpressors, as well as Mp*pif^ko^* plants were grown from gemmae on half-strength Gamborg’s B5 plates with 1% sucrose under cW for 28 days. Mp*DELLA-GR* and *_pro_35S:Cit-*Mp*PIF* Mp*DELLA-GR* lines were grown on sugar-free half-strength Gamborg’s B5 plates under cW for 10 days, then transferred onto plates containing mock solvent or 1 μM DEX and cultured for another 20 days before evaluation. Gemma cups with observable gemmae were observed carefully under stereoscopes, marked on photos taken with a digital camera and then counted. If a gemma with rhizoid and/or growth expansion was observed in a certain gemma cup, it is considered as non-dormant. Representative plants were also photographed with Leica M205C stereo microscopes to show the dormancy of gemma cups in different transgenic lines.

#### Gemma germination assay

Gemma germination assays were carried out following the previous publication^24^. In each experiment, fifty gemmae of each group were planted onto half-strength Gamborg’s B5 plates containing 1% sucrose under green light, then treated with far-red light (30 μmol photons m^−2^ s^−1^) for 15 minutes. For Mp*DELLA-GR* related experiments, 1 μM DEX or the mock solvent was supplemented in the agar plates. After one day of imbibition in the dark, gemmae were irradiated with nothing or a pulse of red light (4500 μmol photons m^−2^) and then cultured for another six days in the dark. Photos of each gemmae were taken using Leica M205C or Olympus SZX16, then evaluated for germination based on growth expansion and/or the development of rhizoids.

#### Gametangiophore formation observation

To monitor the progress of gametangiophore formation in transgenic lines shown in Figure 2B, plants were grown from gemmae on half-strength Gamborg’s B5 plates with 1% sucrose under continuous white light supplemented with far-red light (~20 μmol photons m^−2^ s^−1^, cW+FR). Individual plants were examined and counted for visible gametangiophores each day under stereoscopes. For the experiment in Figure 2I, gemmae of inducible lines or the wild-type control were grown on DEX-free half-strength Gamborg’s B5 plates with 1% sucrose under cW for 10 days, then half pieces of thallus were transferred onto the plates containing 1 nM DEX or mock solvent and cultured under cW+FR. Gametangiophore formation progresses were observed for half plants similarly as described above.

#### Yeast-two hybrid assays

For yeast two-hybrid analyses, Mp*PIF* full length CDS and CDS fragments were amplified from cDNA and introduced into pCR8 using the pCR8/GW/TOPO TA Cloning Kit (Thermo Fisher Scientific) to generate pCR8-MpPIF and -MpPIFdel1-4. Then they were recombined with pGADT7-GW^45^ using Gateway LR Clonase II Enzyme mix (Thermo Fisher Scientific) to produce Gal4-activation domain (AD) fusion proteins. To avoid the previously shown N-terminal transactivation of MpDELLA^7^, only the GRAS domain (pENTR221-MpDELLA^GRAS^) was introduced into pGBKT7-GW^45^ to fuse with the GAL4 DNA-binding domain. Yeast transformation was performed by lithium acetate/single-stranded carrier DNA/polyethylene glycol method as previously described. Y187 and Y2HGold yeast strains were transformed with pGADT7 and pGBKT7-derived expression vectors and selected with Synthetic Defined (SD) medium lacking leucine (-Leu) or tryptophan (-Trp), respectively. Subsequently, haploid yeasts were mated to obtain diploid cells by selection in SD/-Leu-Trp medium. Protein interactions were assayed by the nutritional requirement on histidine (His). SD/-Leu-Trp plates were used as growth control, and SD/-Leu-Trp-His plates supplemented with 5 mM 3-amino-1,2,4-triazole (3-AT, Sigma-Aldrich) was used for interaction evaluation. Spotting assays were performed using cultures with optical density = 1 at a wavelength of 600 nm (OD_600_ = 1) as initial concentration in sequential drop dilutions, and plated with a pin multi-blot replicator. Photos of the same-fold dilutions were taken 3 days after plating.

#### Co-immunoprecipitation (Co-IP) and Bimolecular Fluorescence Complementation (BiFC) assays

Co-IP vectors were obtained by introducing Mp*DELLA* CDS (pENTR221-Mp*DELLA*) into pEarleyGate104 and MpPIF fragments (pCR8-Mp*PIF* and -Mp*PIFdel3*) into pEarleyGate201 ^46^. For BiFC, pENTR211-Mp*DELLA* and pCR8-Mp*PIF* were recombined with pMDC43-YFN and pMDC43-YFC^47^, respectively. *Agrobacterium tumefaciens* GV3101 containing binary plasmids for Co-IP and BIFC were used to infiltrate 4-week-old *Nicotiana benthamiana* leaves.

For Co-IP, leaves were re-infiltrated with a solution of 25 μM MG-132 8 hours before collection 3 days after *A. tumefaciens* infiltration, grinded in liquid nitrogen and homogenized in 1 ml extraction buffer (50 mM Tris–HCl pH 7.5, 150 mM NaCl, 0.1% Triton, 2 mM PMSF, and 1x protease inhibitor cocktail [Roche]). Proteins were quantified using the Bradford assay. 50 μg of total proteins were stored as input. One milligram of total proteins was incubated for 2 h at 4°C with anti-GFP-coated paramagnetic beads and loaded onto μColumns (Miltenyi). Wash and elution from beads was performed according to manufacturer’s instructions. Samples were analyzed by Western-blot after running two 12% SDS-PAGE gels in parallel. One gel was loaded with 25 μg of input, and 10% of eluted proteins; following wet transfer, the PVDF membrane was incubated with an anti-GFP antibody (JL8, 1:5000). The second gel was loaded with 25 μg of input, and 90% of eluted proteins and, after transfer, the membrane incubated with an anti-HA-HRP antibody (3F10, 1:5000). Chemiluminiscence was detected with SuperSignal West Femto substrates (Thermo-Fisher Scientific) and imaged with a LAS-3000 imager (Fujifilm).

For BiFC, leaves were analyzed with a Zeiss LSM 780 confocal microscope. Reconstituted YFP signal was detected with emission filters set to 503-517 nm. Nuclei presence in abaxial epidermal cells was verified by transmitted light.

#### Dual luciferase transactivation assay

Mp*DELLA* and Mp*PIF*-expresssing vectors used for Co-IP (pEarleyGate104-MpDELLA and pEarleyGate201-MpPIF) were used as effector plasmids. A previously available construct with the *Arabidopsis thaliana PIL1* promoter controlling the firefly luciferase gene expression, and a constitutively expressed *Renilla* luciferase gene was used as reporter plasmid^29^. The promoter consists of 1.8 kb upstream of the gene ATG codon, including three consecutive G-boxes known to be bound *in vivo* by PIF3. Transient expression in *N. benthamiana* leaves was carried by agroinfiltration as previously reported^48^. The amount of infiltrated bacteria was set by OD_600_ measurement of A. tumefaciens liquid cultures. Combinations of pre-set reporter-carrying bacteria (OD600 = 0.1) and varying amounts of effector-carrying bacteria were mixed and co-infiltrated together. All the mixes were co-infiltrated alongside *p19* vector-carrying bacteria at a OD_600_ = 0.01. Firefly and the control *Renilla* luciferase activities were assayed in extracts from 1-cm in diameter leaf discs, using the Dual-Glo Luciferase Assay System (Promega) and quantified in a GloMax 96 Microplate Luminometer (Promega). Three leaf disc extracts were quantified per sample in each experiment and repeated for three times. Final quantifications represent means of ratios between firefly luciferase and *Renilla* luciferase read-outs in three independent experiments.

#### RNA isolation, cDNA synthesis, and qRT-PCR analysis

To examine the expression levels of Mp*DELLA* and Mp*PIF* in different transgenic lines, 14-day-old plants grown under cW were homogenized in liquid nitrogen. Total RNA was isolated with TRIzol reagent (Thermo Fisher Scientific) following the manufacturer’s instructions. After checking the concentration and quality of RNA using a NanoDrop 2000 spectrophotometer (Thermo Scientific), up to 3 μg of total RNA was digested with RQ1 RNase-Free DNase (Promega) and reverse-transcribed using ReverTra Ace (Toyobo Life Science). Quantitative real-time PCR (qPCR) was performed in a CFX96 real-time PCR detection system (Bio-Rad) using TaKaRa Ex Taq (TaKaRa Bio) and SYBR Green I Nucleic Acid Gel Stain (Lonza).

For other qPCR experiments, total RNA was extracted with a RNeasy Plant Mini Kit (Qiagen) according to the manufacturer’s instructions. cDNA was prepared from 1 μg of total RNA with PrimeScript 1st Strand cDNA Synthesis Kit (Takara Bio Inc). PCR was performed in a 7500 Fast Real-Time PCR System (Applied Biosystems) with SYBR premix ExTaq (Tli RNaseH Plus) Rox Plus (Takara Bio Inc).

All relative expression levels were calculated following Hellemans *et al.*^49^, and Mp*EF1α* (Mp*ELONGATION FACTOR 1α, Mp3g23400*) was used as the reference gene. Primers are listed in Table S2.

#### RNA sequencing and data analysis

For the Mp*DELLA* overexpression data set, WT and *_pro_35S:*Mp*DELLA-Cit* plants were grown from gemmae on half-strength Gamborg’s B5 plates containing 1% sucrose under LD conditions for 30 days. Then whole plants for two biological replicates were collected for total RNA extraction total RNA with a RNeasy Plant Mini Kit (Qiagen) according to the manufacturer’s instructions. The RNA concentration and integrity [RNA integrity number (RIN)] were measured with an RNA nanochip (Bioanalyzer, Agilent Technologies 2100). Library preparation with TruSeq RNA Sample Prep Kit v.2 (Illumina) and sequencing of 75-nt single-end reads on Illumina NextSeq 550 were carried out at the Genomics Service of the University of Valencia.

For the Mp*pif^ko^* dataset, Tak-1 and Mp*pif^ko^* were grown from gemmae on half-strength Gamborg’s B5 plates containing 1% sucrose under continuous red-light conditions (50 μmol photons m^−2^ s^−1^) for 7 days, then irradiated with far-red light (50 μmol photons m^−2^ s^−1^). Whole plant materials for three biological replicates were collected each at 0, 1, and 4 h after the irradiation. Total RNAs were extracted using RNeasy Plant Mini Kit (Qiagen) and purified with the RNeasy MinElute Cleanup Kit (Qiagen). RNA concentration and qualities were examined by Qubit Assays (Thermo Fisher Scientific) and the Agilent 2100 Bioanalyzer. Libraries were prepared using a TruSeq RNA Sample Prep Kit v.2 (Illumina), quantified by KAPA Library Quantification Kit (Kapa Biosystems), and sequenced of 126-nt single-end reads on Illumina HiSeq 1500 at National Institute for Basic Biology (Okazaki, Japan).

For data processing, reads from both sources were mapped to the *M. polymorpha* reference genome and quantified using Salmon 1.3.0 ^50^. Reads from male lines were mapped to the MpTak1 v5.1 genome^51^, while reads from female Mp*pif^ko^* plants were mapped to autosome sequences from MpTak1 v5.1 plus the known U-chromosome scaffolds from genome ver 3.1 ^11^. Differential gene expressions between sample pairs were analyzed with the R package DESeq2 ^52^, in which both autosome and V chromosome genes were considered for the Mp*DELLA* overexpression set, while only autosome genes were compared between Tak-1 and Mp*pif^ko^*. Genes with a minimum fold change of 2 and adjusted p-value smaller than 0.01 were considered as significantly changed in expression. Compared with the wildtype, *_pro_35S:*Mp*DELLA-Cit* led to the significant up- and down-regulation of 4 and 2 V-chromosome genes, respectively. The total number of MpTak1 v5.1 autosome genes was used for checking if Mp*DELLA*-regulated genes were enriched in differentially expressed genes caused by Mp*pif^ko^* using Fisher’s exact test. UpSet plots were created using the R package ComplexHeatmap^53^.

Fuzzy Gene Ontology (GO)^54^ annotations for the v5.1 (plus ver 3.1 U-chromosome) genes were generated using the Blast2GO algorithm^55^ written in R scripts. Briefly, all *M. polymorpha* reference proteins were blasted^56^ against a database containing all UniProtKB^57^ entries with non-IEA (inferred from electronic annotation) GO annotations, plus all Swiss-Prot entries (release 2020_05) with an e-value threshold of 0.001. Then the GO annotations from top 25 blast hits for each target protein were scored and concatenated based on their similarity and the GO hierarchy (release 2020-06-01). Annotations with scores higher than the user-defined thresholds (40 for cellular component, 55 for biological process, 50 for molecular function) were transferred to *M. polymorpha* proteins. GO enrichment analysis was conducted with biological process terms with the classic fisher’s test from the *topGO* package^58^. R packages *GO.db*^59^, *Rtsne*^60^*, GOSemSim*^61,62^, *rrvgo*^63^ and *AnnotationForge*^64^ were used for the clustering and visualization of top-enriched GO terms. Depositories for the raw sequence datasets, GO annotations and R scripts are listed in the Key Resources Table.

#### Non-targeted flavonoid-related metabolite profiling

Analysis of secondary metabolites in freeze-dried Marchantia samples was carried out following a non-targeted approach as previously described^65^. Briefly, samples (c.a. 10 mg) were extracted in 80% aqueous MeOH containing biochanin A at 1 mg L^−1^ (Sigma-Aldrich, Madrid, Spain) as internal standard by ultrasonication for 10 min. Crude extracts were centrifuged and clean supernatants recovered and filtered through PTFE 0.2 μm syringe filters directly into dark chromatography vials. Extracts were injected into a UPLC system (10 μL) (Acquity SDS, Waters Corp. Ltd. USA) and separations carried out on a C18 column (Luna Omega Polar, C18, 1.6 μm, 100 × 2.1 mm, Phenomenex, CA, USA) using acetonitrile and ultrapure water, both supplemented with formic acid at a concentration of 0.1% (v/v), as solvents at a flow rate of 0.3 mL min^−1^. A gradient elution program starting from 5% to 95% acetonitrile in 17 min followed by a 3 min re-equilibration period was employed. Compounds were detected by mass spectrometry using a hybrid quadrupole time-of-flight mass spectrometer (QTOF-MS, Micromass Ltd., UK) coupled to the UPLC system through an electrospray source. Samples were analyzed in both positive and negative electrospray modes within 50-1000 Da mass range using two simultaneous acquisition modes: 1) low CID energy for profiling purposes and 2) high CID energy for MS/MS of selected compounds, this was achieved by setting an energy ramp from 5-60 eV. During measurements cone and capillary voltages were set at 30 V and 3.5 kV, respectively; source and block temperatures were kept at 120°C. Desolvation gas (N_2_) was kept at 350 °C at a flow rate of 800 L h^−1^. Nebulization gas was also N_2_ at a flow rate of 60 L h^−1^. In the collision cell, pure Ar was used as the collision gas. Exact mass measurements were achieved by monitoring the reference compound lockmass leucine-enkephalin ([M+H]^+^ 556.2771 and [M-H]− 554.2514, respectively); therefore, the resulting mass chromatograms were acquired in centroid mode.

Processing of mass chromatograms was performed with xcms^66^ after conversion to mzXML with MSConvert^67^ using default settings. Chromatographic peak detection was performed using the matchedFilter algorithm, applying the following parameter settings: snr = 3, fwhm = 15 s, step = 0.01 D, mzdiff = 0.1 D, and profmethod = bin. Retention time correction was achieved in three iterations applying the parameter settings minfrac = 1, bw = 30 s, mzwid = 0.05D, span = 1, and missing = extra = 1 for the first iteration; minfrac = 1, bw = 10 s, mzwid = 0.05 D, span = 0.6, and missing = extra = 0 for the second iteration; and minfrac = 1, bw = 5 s, mzwid = 0.05 D, span = 0.5, and missing = extra = 0 for the third iteration. After final peak grouping (minfrac = 1, bw = 5 s) and filling in of missing features using the fillPeaks routine of the xcms package, a data matrix consisting of feature × sample was obtained. When available, identification of metabolites was achieved by comparison of mass spectra and retention time with those of authentic standards or alternatively were tentatively annotated by matching experimental mass spectra in public databases (metlin, Massbank or HMDB). Known and temptative flavonoid-related compounds were chosen for comparison. Before statistical analyses, raw peak area values were normalized to internal standard area and sample weight. Pairwise comparisons (wild type vs *_pro_35S*:Mp*DELLA-Cit*) were carried out using a two-tailed Student’s t-test comparing two groups of samples of identical variance.

#### Analysis of survival after oxidative stress

For survival measurement, 10 gemmae per genotype and experiment were grown on top of Whatman filter papers discs (Thermo Fisher) on half-strength Gamborg’s B5 1% agar medium for 6 days, and then transferred to half-strength Gamborg’s B5 1% agar medium supplemented with 100 μM methyl viologen to produce a severe oxidative stress for 10 days. Gemmallings were transferred back to half-strength Gamborg’s B5 1% agar medium for recovery. Survival was counted when independent apical regions resumed growth and represented as the percentage of growth-resuming apical regions of the total number at the beginning of the stress treatment.

In Mp*DELLA-GR* related assays, the same procedure was followed, but mock and 1 μM dexamethasone (DEX) were included for DELLA activity induction during the 10 days of oxidative stress treatment. In addition, DEX or mock (ethanol) were added in water solution 24 hours before stress treatment.

## Supplemental Information

Supplementary figures 1-4.

Table S1. Differentially accumulated flavonoid-related metabolites in Marchantia polymorpha wild-type and *pro35S:*Mp*DELLA-Cit* line. Related to Figure S4C.

Table S2. List of oligonucleotides, related to the STAR Methods.

Data S1. RNA-seq expression profiles, GO annotations and top-enriched GO terms. Related to Figure 3.

