## Supplementary material for "Coordination between growth and stress responses by DELLA in the liverwort *Marchantia polymorpha*": Supplemetary Figures and Tables

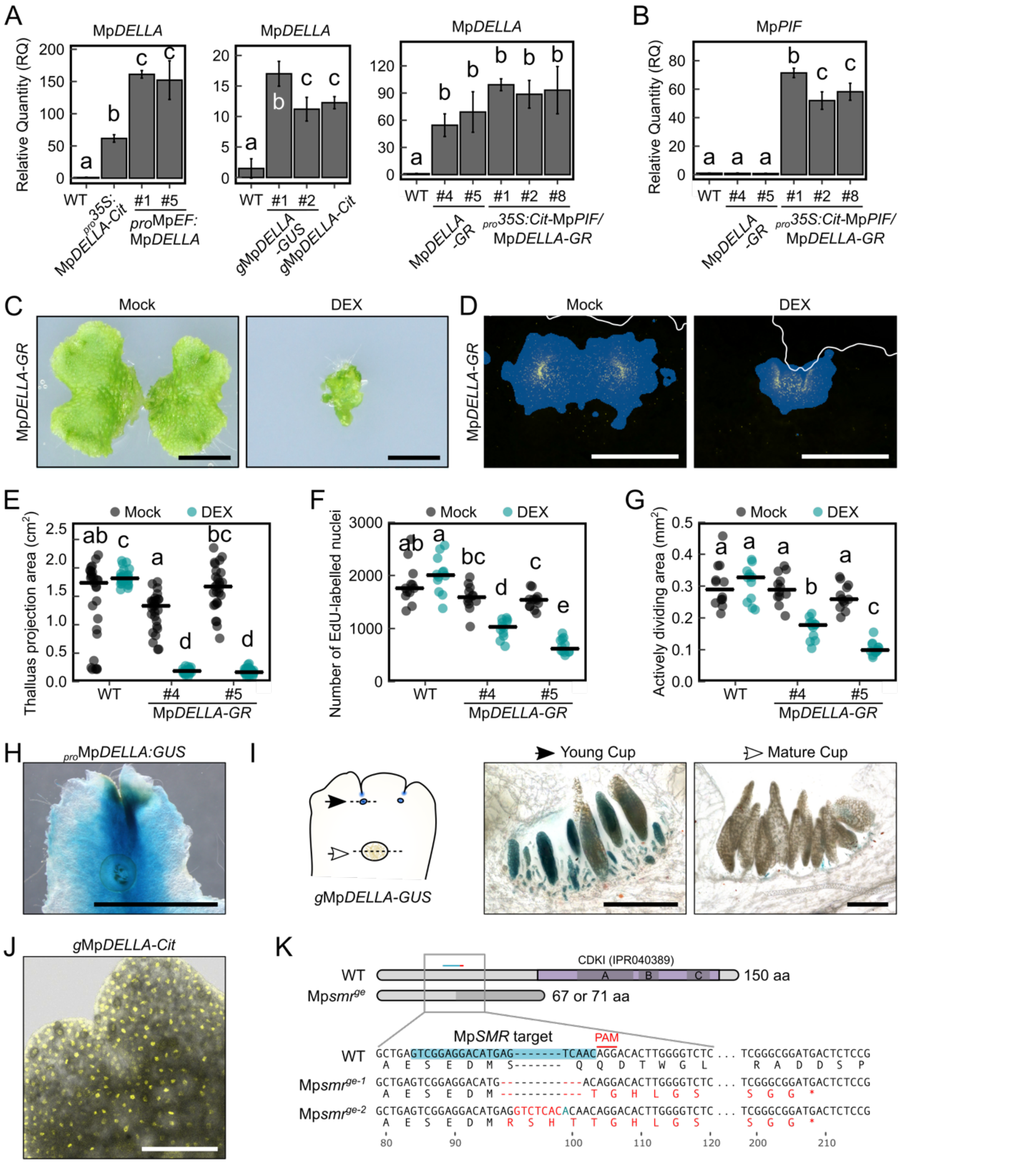


**Figure S1. Related to Figure 1.**

(A) Relative expression level of Mp*DELLA* determined by RT-qPCR in 14-day-old gemmalings of multiple transgenic lines.

(B) Relative expression level of Mp*PIF* determined by RT-qPCR in 14-day-old gemmalings in $_{pro}$*35S:*Mp*PIF-Cit* Mp*DELLA-GR* lines

(C) Morphology of Mp*DELLA-GR* plants, grown from gemmae for 14 days with mock or 1 μM DEX. Scale bar, 5 mm.

(D) Apical notches of 7-day-old Mp*DELLA-GR* gemmalings, treated with mock or 1 μM DEX for two days and labelled with EdU (yellow signals). Plant boundaries are marked with white lines, and blue color indicates the area occupied by dividing cells. Scale bar, 500 μm.

(E) Measurement of plant sizes in Mp*DELLA-GR* gemmalings, grown from gemmae with mock or 1 μM DEX for 14 days. n=27.

(F) Number of EdU-labelled nuclei in the apical notches of 7-day-old Mp*DELLA-GR* gemmalings, treated with mock or 1 μM DEX for two days. n=12.

(G) Quantification of actively dividing area in the apical notches of 7-day-old Mp*DELLA-GR* gemmalings, treated with mock or 1 μM DEX for two days. n=12.

(H) GUS-stained thallus of 21-day-old $_{pro}$Mp*DELLA:GUS*. Scale bar, 5 mm.

(I) Agar sectioning of developing (black arrow) and mature (white arrow) gemma cups of GUS-stained thallus of 21-day-old *g*Mp*DELLA-GUS*. Scale bars, 200 μm.

(J) Microscopic image of the apical notch of *g*Mp*DELLA-Cit* gemma showing MpDELLA-Cit nuclear localization. Scale bar, 200 μm.

(K) Genotype information for the Mp*smr*$^{ge}$ Mp*DELLA-GR* lines. Predicted protein products from wild-type and genome-edited Mp*SMR* locus were illustrated (purple boxes: CDKI functional domains; dark-grey shade: frameshifts caused by genome editing). Sequences of wild-type and both CRISPR/Cas9-derived alleles were shown in alignments. The guide RNA target sequence is highlighted in blue. Deletions and insertions were indicated with red letters and the substitution with blue ones.

In A, B, error bars represent standard error of three biological replicates. In E, F, G, dots represent individual plants and horizontal lines represent mean values. Statistical groups were determined by Tukey's Post-Hoc test ($p<0.05$) following one-way ANOVA.

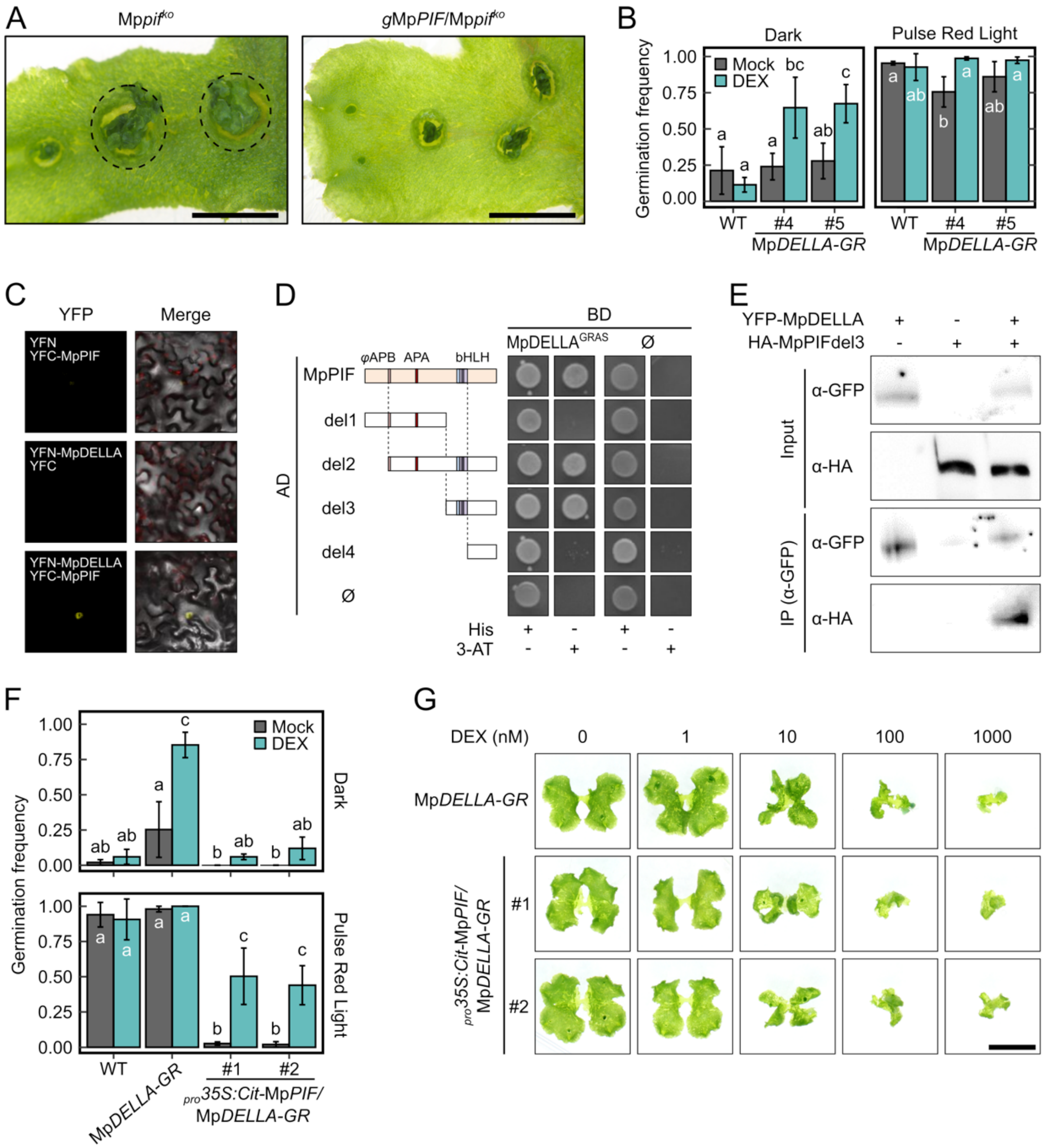


**Figure S2. Related to Figure 2.**

(A) Images of 28-day-old plants showing premature gemmae germination inside the cups of Mp*pif*$^{ko}$ and the complemented line (*g*Mp*PIF*/Mp*pif*$^{ko}$). Dashed circles indicate non-dormant gemma cups. Scale bar, 5 mm.

(B) Germination frequencies of the wild-type and Mp*DELLA-GR* gemmae under different light conditions. Gemmae were imbibed and treated without (Dark) or with a pulse of red light (4500 μmol photons m$^{-2}$) followed by incubation in the dark. Dark grey and turquoise columns represent gemmae supplemented with mock or 1 μM DEX, respectively.

(C) Bimolecular fluorescence complementation assays showing MpDELLA-MpPIF interaction in *N. benthamiana* abaxial leaves.

(D) Physical interaction between MpDELLA GRAS domain and MpPIF bHLH domain shown by yeast two-hybrid assays of MpPIF deletion fragments. Although there is no conserved APB domain in MpPIF, its theoretical position was marked by *ψ*APB and used for fragmentation. MpPIF amino acids present on each fragment are 1-472 (del1), 132-760 (del2), 473-760 (del3) and 588-760 (del4). Histidine (His) supplemented media used as growth control. 5 mM 3-amino-1,2,4-triazole (3-AT) was added to His-depleted medium.

(E) Physical interaction between YFP-MpDELLA and HA-MpPIFdel3 (aa 473-760) shown by co-immunoprecipitation after agroinfiltration in *N. benthamiana* leaves.

(F) Germination frequencies of Mp*DELLA-GR* gemmae with additional expression of Mp*PIF* ($_{pro}$*35S:Cit*-Mp*PIF*) under different light conditions as indicated in (B). Dark grey and blue columns represent gemmae supplemented with mock or with 1 µM DEX, respectively.

(G) Morphology of 14-day-old Mp*DELLA-GR* and $_{pro}$*35S:Cit*-Mp*PIF/*Mp*DELLA-GR* plants, grown with different concentrations of DEX. Scale bar, 1 cm.

In B, F, error bars represent standard deviation from three independent experiments (n = 50 per experiment). Statistical groups were determined by Tukey's Post-Hoc test ($p<0.05$) following ANOVA analysis.

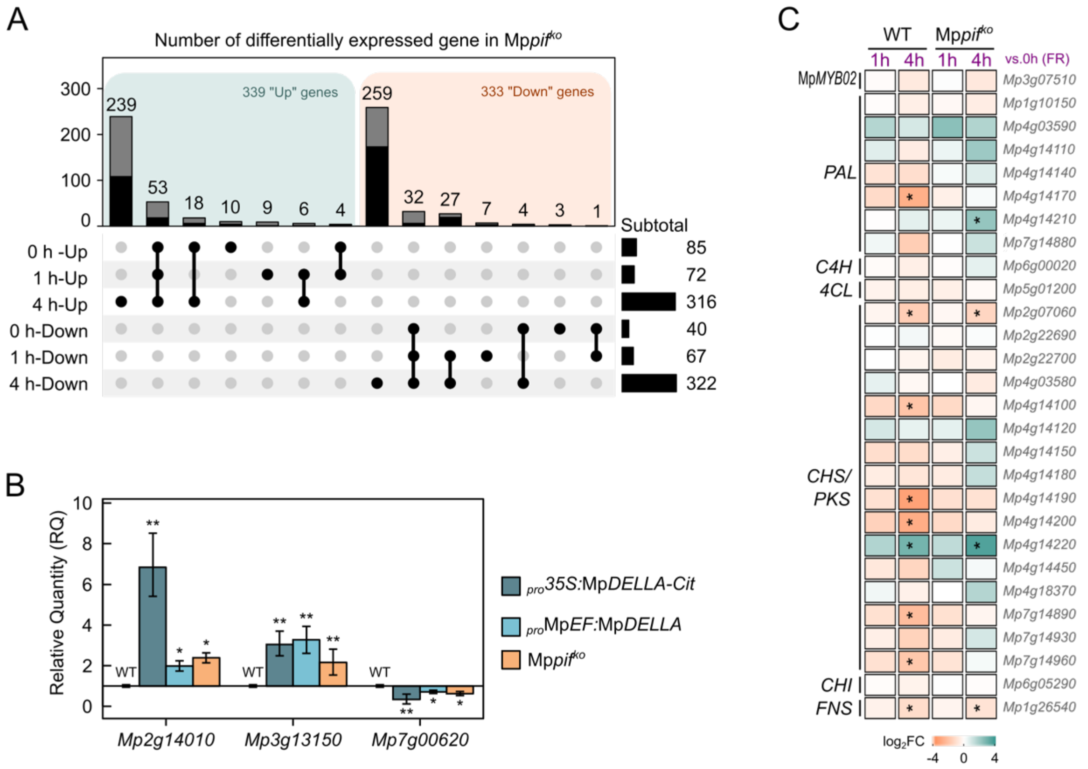


**Figure S3. Related to Figure 3.**

(A) UpSet plot showing differentially expressed genes between Mp$pif^{ko}$ and the wild-type at different time points after far-red light (FR) irradiation. Black proportions in the top column plot represent genes ever changed significantly (with |log2FC|>1 and adjusted p<0.01 calculated by DESeq2) in response to far-red treatment in either genotypes.

(B) Relative expression level of selected genes by RT-qPCR, as a verification for the RNA-seqs. Error bars represent standard deviation from three biological replicates. Asterisks indicate statistically significant differences with respect to the wild type (*, p<0.05; **, p<0.01, after a Student's t-test)

(C) Fold changes in expression (log2 scale) of genes related to flavonoid biosynthesis, compared to time point 0 in both genotypes. Asterisks indicate genes considered as significantly changed (with |log2FC|>1 and adjusted p<0.01 calculated by DESeq2).

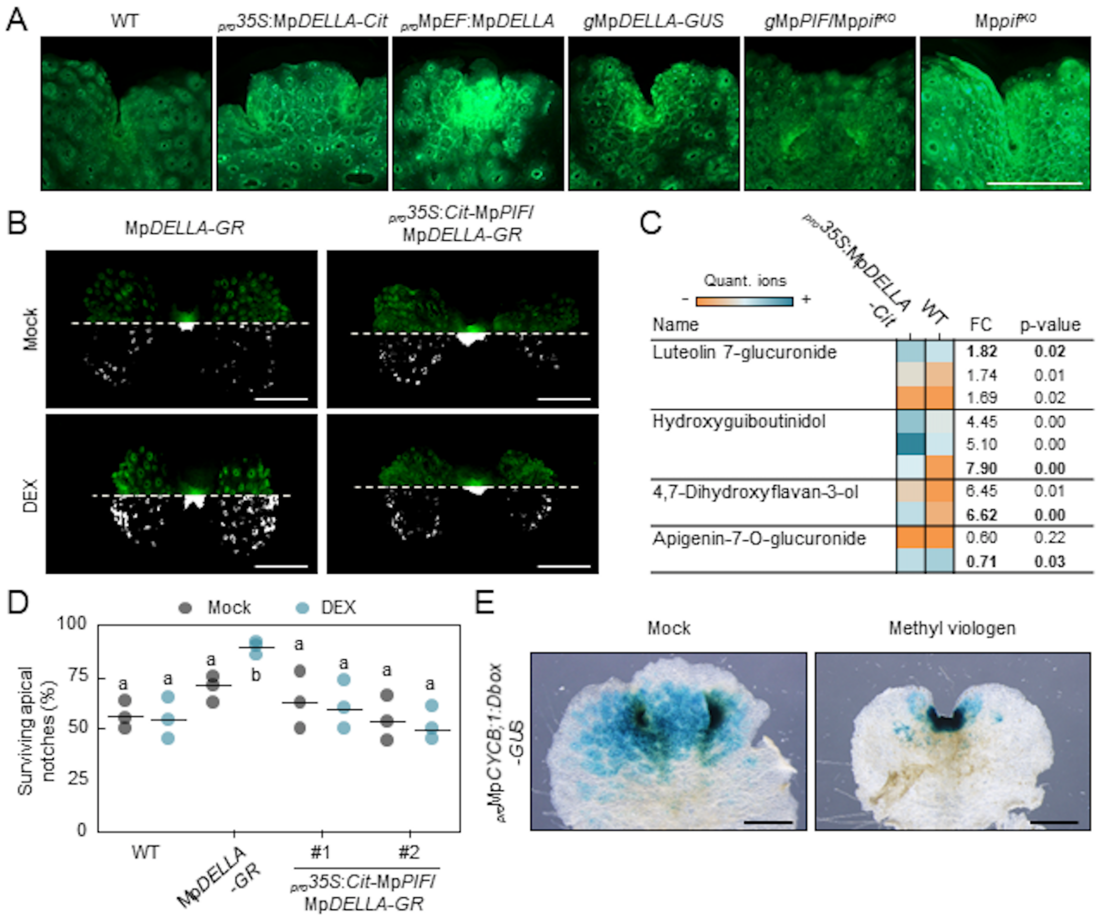


**Figure S4. Related to Figure 4.**

(A) Representative images of apical region of Figure 4A gemmalings, stained by DPBA. More intense fluorescent signals denote higher general flavonoid content. Scale bar, 1 mm.

(B) Representative images showing DPBA staining of 9-day-old Mp*DELLA-GR* and $_{pro}$*35S:Cit*-Mp*PIF* Mp*DELLA-GR* gemmalings, grown with or without 1 μM DEX for 3 days. Upper halves, original picture taken with GFP filter; Lower halves, pseudo-color intensity binary maps (threshold at 30%) to facilitate the comparison of the differences in fluorescence signal between images. Scale bar, 2 mm.

(C) Differentially accumulated flavonoid-related compounds in wild-type and $_{pro}$*35S*:Mp*DELLA-Cit*. Quantifier ions are listed in bold letters. FC, fold change of overexpressor versus wild-type. Original data with individually detected ions and parameters of detection can be found in Table S1.

(D) Percentage of surviving apical notches after a 10-day treatment with 100 μM methylviologen (MV). Mp*DELLA-GR* induction with 1 µM DEX started one day earlier before MV application and was further maintained throughout the stress treatment.

(E) Images of 13-day-old Mp*CYCB;1-GUS* Mp*DELLA-GR* plants stained for GUS activity after mock or 10 µM MV treatment for 6 days. Scale bar, 1 mm.

In B, fold changes and p-values from Student's t-tests are calculated with quantifications from four samples per genotype. In D, E, dots represent biological replicates and horizontal lines median values from three independent experiments, statistical support is provided by one-way ANOVA analysis, and groups determined by Tukey's Post-Hoc test ($p<0.01$).

**Table S1. Differentially accumulated flavonoid-related metabolites in *Marchantia polymorpha* wild-type and *$_{pro}$35S:*Mp*DELLA-Cit* line. Related to Figure S4C.**

| Name | mz | rt | adduct | Tak-1_1 | Tak-1_2 | *$_{pro}$35S:MpDELLA-Cit_1* | *$_{pro}$35S:MpDELLA-Cit_2* | FC | p value (t-test) |
|---|---|---|---|---|---|---|---|---|---|
| Luteolin 7-O-glucuronide | 287.0578 | 435.1638 | $[M+H-Hexose]^+$ | 337.19652 | 162.45369 | 410.06821 | 479.18282 | 1.78 | 0.17 |
| | 463.0875 | 435.6399 | $[M+H]^+$ | 22340.093 | 17792.167 | 36658.479 | 36374.806 | 1.82 | 0.02 |
| | 464.0916 | 435.2707 | | 5391.2870 | 4490.9237 | 8619.7590 | 8585.8281 | 1.74 | 0.01 |
| | 465.0935 | 434.6752 | | 1155.6397 | 1128.9224 | 2047.7415 | 1821.4503 | 1.69 | 0.02 |
| | 46.6095 | 434.6273 | | 157.58420 | 212.97500 | 87.648122 | 448.65807 | 1.45 | 0.69 |
| | 485.0645 | 434.6398 | $[M+Na]^+$ | 155.28937 | 220.78331 | 187.79761 | 159.25366 | 0.92 | 0.72 |
| Luteolin 7,3'-di-O-glucuronide | 639.1187 | 370.7217 | $[M+Na]^+$ | 14625.707 | 10749.556 | 28421.449 | 27464.365 | 2.2 | 0.02 |
| | 640.1209 | 370.7217 | | 4082.7649 | 2912.6274 | 8007.6455 | 8245.0167 | 2.32 | 0.02 |
| Hydroxyguiboutinidol (tentative) | 107.0515 | 339.3267 | | 11086.915 | 8490.3717 | 43589.294 | 43506.816 | 4.45 | 0 |
| | 153.0573 | 338.8445 | | 2542.3915 | 1341.2988 | 6100.9561 | 6169.3699 | 3.16 | 0.02 |
| | 165.0568 | 339.3267 | $[M+H-C_6H_8O_2]^+$ | 2957.3126 | 2218.8410 | 11953.959 | 11978.778 | 4.62 | 0 |
| | 259.0983 | 339.3865 | $[M+H-H_2O]^+$ | 19013.674 | 15643.135 | 87566.923 | 89226.251 | 5.1 | 0 |
| | 277.1099 | 339.4859 | $[M+H]^+$ | 1709.4595 | 1232.7364 | 11915.684 | 11319.681 | 7.9 | 0 |
| 4,7-Dihydroxyflavan-3-ol | 121.0678 | 578.9766 | $[M+H-CO_2]^+$ | 342.99611 | 408.00936 | 1474.9217 | 1096.5793 | 3.42 | 0.04 |
| | 14.7046 | 576.6364 | $[M+H-H_2O]^+$ | 1102.3484 | 1084.8219 | 7550.9103 | 6561.5044 | 6.45 | 0.01 |
| | 165.0569 | 577.3081 | $[M+H-C_6H_8O_2]^+$ | 284.67468 | 411.38642 | 1839.6545 | 1796.7498 | 5.22 | 0 |
| | 241.0879 | 576.6364 | $[M+H-H_2O]^+$ | 3728.6684 | 3480.0925 | 25187.633 | 22516.680 | 6.62 | 0 |
| Apigenin-7-O-glucuronide | 271.0633 | 424.9005 | $[M+H-C_6H_8O_6]^+$ | 4251.8180 | 3806.1232 | 3308.3657 | 3088.4613 | 0.79 | 0.08 |
| | 272.0669 | 424.3484 | | 445.44160 | 708.27758 | 345.49883 | 346.97561 | 0.6 | 0.22 |
| | 447.0928 | 424.8175 | $[M+H]^+$ | 36104.524 | 32573.717 | 24223.723 | 24750.314 | 0.71 | 0.03 |
| | 448.0967 | 424.4458 | | 8525.8855 | 7470.6678 | 5410.8245 | 6273.9972 | 0.73 | 0.09 |
| | 449.0973 | 424.8175 | | 1744.8665 | 1584.7019 | 1228.1131 | 1409.2066 | 0.79 | 0.1 |

**Table S2. List of oligonucleotides, related to the STAR Methods**

| Name | Sequence (5' to 3') | Used for |
|---|---|---|
| MpDELLA_Fw2 | GGGGACAAGTTTGTACAAAAAAGCAGGCTTAATGGATTCCTCTGCCGATTACG | pENTR221-MpDELLA |
| MpM5DELLA_Fw2 | GGGGACAAGTTTGTACAAAAAAGCAGGCTACATCTCGGATTCAATGGCTGGAG | pENTR221-MpΔDELLA |
| pMpDELLA_Fw2 | GGGGACAAGTTTGTACAAAAAAGCAGGCTCGGCGTAGGAGATGGGCACTTG | pENTR221-$_{pro}$MpDELLA<br>pENTR221-gMpDELLA |
| pMpDELLA_Rv2 | GGGGACCACTTTGTACAAGAAAGCTGGGTAACAGCAGGCAATTATCACTCTTCG | pENTR221-$_{pro}$MpDELLA |
| MpDELLA_Rv2 | GGGGACCACTTTGTACAAGAAAGCTGGGTAGGAACAATGCCATGCCGATG | pENTR221-MpDELLA,<br>pENTR221-MpΔDELLA,<br>pENTR221-gMpDELLA |
| CACC-MpDELLA-CDS-F | CACCATGGATTCCTCTGCCG | pENTR-MpDELLA |
| MpDELLA-CDS-ns-R | GGAACAATGCCATGCCGATG | pENTR-MpDELLA |
| SalI-MpDELLApro-F | CGCGTCGACATAGAATACGCAACTTTATGGCA | pENTR1A-$_{pro}$MpDELLA-short |
| NotI-MpDELLApro-R | AAAGCGGCCGCTAACAGCAGGCAATTATCACTCT | pENTR1A-$_{pro}$MpDELLA-short |
| MpDELLApro1-IF-F | AGGAACCAATTCAGTCGACACGGCGTAGGAGATGGG | pENTR1A-$_{pro}$MpDELLA,<br>pENTR1A-gMpDELLA |
| MpDELLApro2-IF-R | CCTTTTGCCATAAAGTTGCGTATT | pENTR1A-$_{pro}$MpDELLA,<br>pENTR1A-gMpDELLA |
| MpDELLA-cassette-IF-F | AATTGCCTGCTGTTAATGGATTCCTCTGCCGATTACG | pENTR1A-gMpDELLA-shhort |
| MpDELLA-cassette-IF-R1 | ATATCTCGAGTGCGGGGAACAATGCCATGCCGATG | pENTR1A-gMpDELLA-short |
| SalI-MpCYCB-5' * | GTCGACTCAAAAATTCTCCTCCGTACA | pENTR1A-MpCYCB;1-Dbox |
| MpCYCB-116-3'-pENTR | GCCCCGAAGCAGGAGCAATGT | pENTR1A-MpCYCB;1-Dbox |
| MpPIF_1F | CACCATGAGTCACCTCGTTC | pENTR-MpPIF-Stop |
| MpPIF_0R | AAGCAAGCGTGGAAATCAAG | pENTR-MpPIF-Stop |
| MpPIF_Fw2 | ATGAGTCACCTCGTTCCCG | pCR8-MpPIF<br>pCR8-MpPIFdel1 |
| MpPIF_Rv2 | TTGGGGGGCTCCACCGCCC | pCR8-MpPIF<br>pCR8-MpPIFdel2-4 |
| MpPIFdel1_Rv | GGCCTCTTTCCCTCTATC | pCR8-MpPIFdel1 |
| MpPIFdel2_Fw | CAGGAAGACGAGATGGTG | pCR8-MpPIFdel2 |
| MpPIFdel3_Fw | GCATCTAGTGGCAAGAGA | pCR8-MpPIFdel3 |
| MpPIFdel4_Fw | CAGATGATGTCCATGAGA | pCR8-MpPIFdel4 |
| sgRNA_MpSMR_F | CTCGGTCGGAGGACATGAGTCAAC | pMpGE_En03-MpSMR-gRNA |
| sgRNA_MpSMR_R | AAACGTTGACTCATGTCCTCCGAC | pMpGE_En03-MpSMR-gRNA |
| genoMpSMR_F | GGTAGTTCCTCTGGCTCAAG | MpSMR genotyping |
| genoMpSMR_R | CTGTGATACGGTAGGAATGAGT | MpSMR genotyping |
| MpEF1−qPCR_F | AAGCCGTCGAAAAGAAGGAG | qPCR (Mp3g23400) |
| MpEF1−qPCR_R | TTCAGGATCGTCCGTTATCC | qPCR (Mp3g23400) |
| MpDELLA-RT-F1 | AGTTCTACGAGACTTGTC | qPCR (Mp5g20660) |
| MpDELLA-RT-R1 | ATGTATCCGCTTGTGATT | qPCR (Mp5g20660) |
| MpPIF-qPCR-F1 | CAGCCGATGAGTATGGATGC | qPCR (Mp3g17350) |
| MpPIF-qPCR-R1 | AGATGATGGAGCGAATGCTG | qPCR (Mp3g17350) |
| MpPALd-qPCR-F | CTGCTAAGAAATCTCTATTCAC | qPCR (Mp7g14880) |

| | | |
|---|---|---|
| MpPALd-qPCR-R | ACTAGTGGCATCATCTATGTAA | qPCR (Mp7g14880) |
| MpPALb-qPCR-F | CACTTACGGTGTCACTACAG | qPCR (Mp4g14110) |
| MpPALb-qPCR-R | ATCAACTCTCTCTGTAAGTCG | qPCR (Mp4g14110) |
| MpC4Ha-qPCR-F | GAGAAAGCAGCTATTGATTAC | qPCR (Mp8g00020) |
| MpC4Ha-qPCR-R | GTAGTCTCAATAGCAGCAAC | qPCR (Mp8g00020) |
| MpCHS/STBS-qPCR-F | AGTTCGCTCGAATTTGTAAG | qPCR (Mp4g18370) |
| MpCHS/STBS-qPCR-R | AAGTGAGGGATCCTTGTAC | qPCR (Mp4g18370) |
| MpGH3B-qPCR-F | GTAAACTGAAAATGCTACCTGT | qPCR (Mp2g14010) |
| MpGH3B-qPCR-R | GTTGATTTCATGTGAAACC | qPCR (Mp2g14010) |
| MpKSb-qPCR-F | GACCTTCCGTACAATCTC | qPCR (Mp3g13150) |
| MpKSb-qPCR-R | GATGAAGGGGTATTTTGC | qPCR (Mp3g13150) |
| MpFUT-qPCR-F | TGACTTCACATATGGTTATACC | qPCR (Mp7g00620) |
| MpFUT-qPCR-R | CATAATCGTATAAGCTCTTGAC | qPCR (Mp7g00620) |

* The last nucleotide of this primer does not match with the current reference genome, though the amplification was successful and confirmed by sequencing.
Additional overhangs for proper cloning (Gateway/restriction site/sticky end) are marked in red; start codons are underlined.
